# Performant web-based interactive visualization tool for spatially-resolved transcriptomics experiments

**DOI:** 10.1101/2023.01.28.525943

**Authors:** Chaichontat Sriworarat, Annie Nguyen, Nicholas J. Eagles, Leonardo Collado-Torres, Keri Martinowich, Kristen R. Maynard, Stephanie C. Hicks

## Abstract

High-resolution and multiplexed imaging techniques are giving us an increasingly detailed observation of a biological system. However, sharing, exploring, and customizing the visualization of large multidimensional images can be a challenge. Here, we introduce Samui, a performant and interactive image visualization tool that runs completely in the web browser. Samui is specifically designed for fast image visualization and annotation and enables users to browse through large images and their selected features within seconds of receiving a link. We demonstrate the broad utility of Samui with images generated with two platforms: Vizgen MERFISH and 10x Genomics Visium Spatial Gene Expression. Samui along with example datasets is available at https://samuibrowser.com.

## Background

Recent technological advances have led to the generation of increasingly large and multi-dimensional images that can be used for spatially-resolved transcriptomics (SRT) [1]. This enables researchers to map transcriptome-wide gene expression data to spatial coordinates within intact tissue at near- or sub-cellular resolution. By combining two distinct data modalities, this technology can yield unprecedented insight into the spatial regulation of gene expression [2]. Yet, this combination poses a unique computational and visualization challenge – we now have high-resolution images with tens of thousands of dimensions of features, each corresponding to a gene from the transcriptome.

Existing open-source image processing tools, such as ImageJ / Fiji [3] and napari [4] are often used for image visualization, but are not designed to view more than a few dimensions of data. Large images are commonly distributed using the TIFF (Tag Image File Format). To browse and interact with these datasets, users need to load the entire dataset into memory, which limits the use of these datasets to specialized workstations with large amounts of memory (e.g. tens to hundreds of gigabytes) [5]. This computational requirement limits the larger scientific community from accessing many datasets. Ability to access these datasets is also limited by the need to download datasets prior to visualization.

An alternative solution are web-based visualization tools, which allow users to view images without downloading. For example, neuroglancer [6], a volumetric data viewer, allows users to view high-dimensional datasets with a hyperlink. However, its original design goal as a connectomics viewer means that it cannot view high-dimensional annotations. Interactive web-based applications in R, such as Shiny apps [7], are a common vehicle but they rely on a server instance of R, which limits scalability and unavoidably introduces latency.

Here, we developed a web-based open source application, Samui. Built on top of a GPU-accelerated geospatial analysis package, OpenLayers [8], Samui can responsively display large, multi-channel images (>20,000 × 20,000 pixels) and their annotations in the web browser. In addition to local files, Samui can directly open processed images from a cloud storage link, such as Amazon Web Services (AWS) S3 storage, without the need to download the entire dataset.

To achieve this level of performance, Samui utilizes the cloud-optimized GeoTIFF (COG) file format [9], which is designed by the Geographic Information Systems (GIS) community to display raster maps. This format has the ability to organize the full image into tiles of varying resolution, allowing Samui to only selectively load sections of the image that the user is viewing. Feature data, such as gene expression or cell segmentation, are also similarly chunked and loaded on demand. This ‘lazy’ technique drastically reduces the download requirements and memory consumption of Samui.

## Results and Discussion

Samui enables interactive and responsive visualization of large, multidimensional images that can easily be shared with the larger scientific community (**Figure 1**). We focus our efforts here to provide an overview of the Samui, and describe key innovations compared to currently available open-source image visualization tools with examples from SRT including spatial barcoding followed by sequencing [10–13] or fluorescence imaging-based transcriptomics [14, 15].

**Figure 1:**
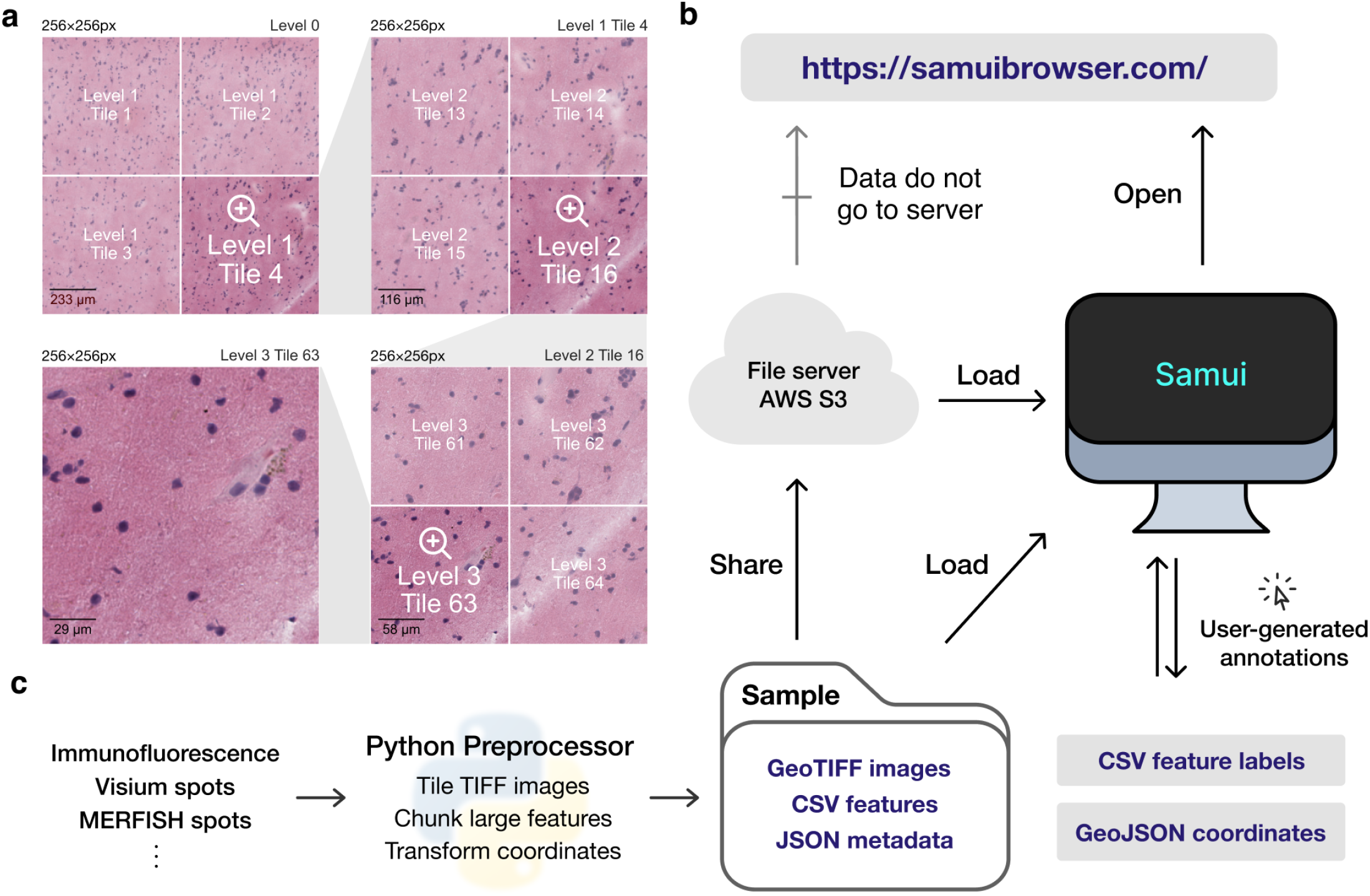
Overview of the Samui. **(a)** Input images are stored in a tiled GeoTIFF file format. When a user opens the browser, at the lowest zoom setting, only tiles from level 0 are downloaded. With increasing Zoom setting, tiles from increasing levels within the field of view are downloaded as needed. **(b)** Samui is statically served and relies completely on the client machine for all functionalities. It can load files from a cloud storage link or from the local machine. (**c**) Samui takes in a Sample folder that contains chunked images, CSV features, and JSON metadata. These are generated with the Python preprocessor and the browser also supports user-generated annotations.

The browser primarily takes as input (i) a JSON (Javascript object notation) file that describes the metadata of the dataset, (ii) a Cloud-optimized GeoTIFF (COG) file containing *n*-dimensional tiled images as pixel intensity (**Figure 1a**). Feature information (e.g. gene expression data) can be represented using standard CSV (comma-separated value) files, but can also be represented by a cluster of sparse CSVs that has been chunked and compressed for fast readout. COG was chosen over alternative chunked file formats, such as HDF5 [16] and Zarr [17] file formats, as the byte-indexable structure of COGs easily enables simple HTTP GET [18] range requests can be used to retrieve fine-grained portions of the COG. It also is specialized for image data and is rigorously specified by the OGC GeoTiff Standard [19].

Samui is statically served and relies completely on the client machine for all functionalities. Users can open these files directly from their local file system. In addition, as a web application, Samui can directly open the links to the hosted images in cloud storage without the need to fully download them first (**Figure 1b**). This approach of web-based visualization has been successfully applied for other data types including genomics [20], geospatial [21], and astronomy [22].

The input files can be generated using a provided Python-based command line tool and GUI (graphical user interface) provided along with Samui. To support SRT data, Samui offers functionality to convert standard output from quantification tools such as 10x Genomics Space Ranger to these cloud-optimized file formats (**Figure 1c**).

As noted above, a particularly important data analysis step of SRT bio-images is annotation. Samui can be used to rapidly annotate features, such as Visium spots (**Figure 2a**) or individual cell types (**Figure 2b**), that can be exported as a CSV file. It can also import existing annotations from other software, such as from napari or ImageJ/Fiji. For example, using data from the 10x Genomics Visium Spatial Gene Expression platform, a Samui user can annotate at the cell (segmented cells/points on the image), spot (55 μm circles), or spatial domain (polygons) level (**Figure 2a**). Annotation can also be done using data acquired from the 10x Genomics Visium Spatial Proteogenomics platform, which generates multiplex images of fluorescently-labeled proteins paired with gene expression [23].

**Figure 2:**
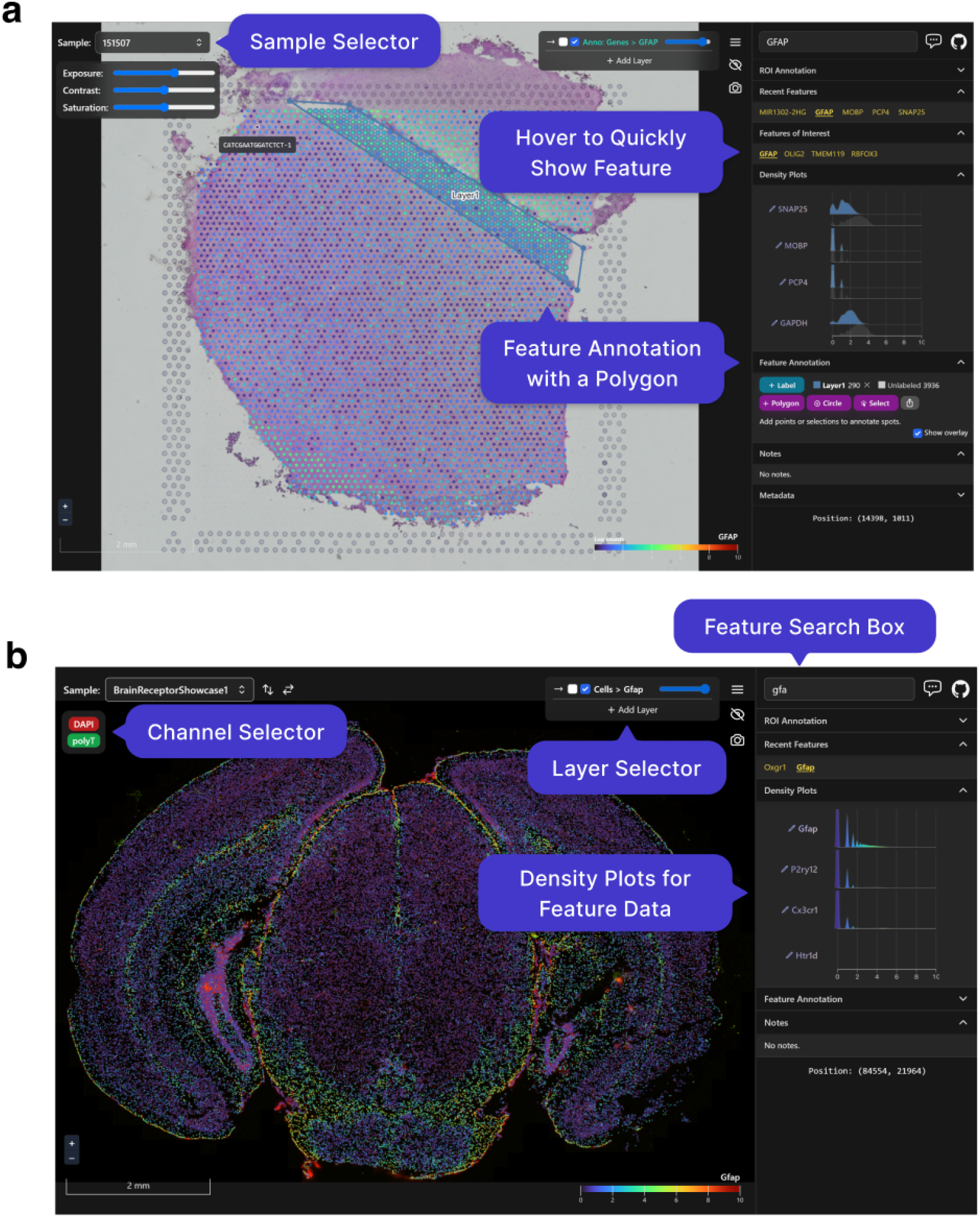
Visualizing and annotating multi-dimensional images in Samui. A user can easily interact with and annotate spatial coordinates on top of multidimensional images with data such as from the **(a)** 10x Genomics Visium Spatial Gene Expression platform or **(b)** Vizgen MERFISH platform.

## Conclusions

Here, we developed Samui, an open source and web-based image visualization system. We demonstrated the broad utility of the Browser with a rapid data exploration of images generated with Vizgen MERFISH and the 10x Genomics Visium Spatial Gene Expression or Proteogenomics platforms. The key strengths of the Samui compared with existing open-source image tools [3, 4, 24, 25] for visualization and annotation include (i) leveraging cloud-optimized GeoTIFF files that efficiently stream large scientific images, (ii) leveraging web-based browser to easily visualize and share images with the scientific community, and (iii) the ability to easily annotate images, which is particularly important in SRT applications. Our visualization tool can be used to view multiple modalities at the same time along with multiple images each with individual *n*-dimensional information. The features offered in Samui complement those of other visualization tools and provide researchers with a better understanding of these complex datasets.

## Methods

### Samui

Samui is written with SvelteKit v1.0 and TypeScript v4.6. The latest version of Samui is available at https://github.com/chaichontat/samui.

The WebGL image renderer is an adaptation from OpenLayers v6.15.1, a geospatial visualization framework. Samui displays images with the cloud-optimized GeoTIFF (COG) format and stores other data in the CSV format. Each dataset is organized into a “Sample” folder that contains a master JSON (Javascript object notation) file with the metadata of the dataset. This file provides links to other COGs and CSVs that describe different overlays and features.

As data in spatial transcriptomics tend to share a common coordinate, we separated the notion of coordinates and features in Samui to reduce redundancy. An overlay can be thought of as a layer of points, each with a spatial coordinate. Features are the categorical or quantitative values associated with each point in an overlay. For example, in a Visium experiment, the position of a Visium spot would be stored as an overlay whereas the expression of a gene at each spot would be a feature. For large datasets, CSVs can be sparsified, chunked, and compressed to allow lazy loading of feature data. This is done in the Visium and MERFISH example described below.

### Input data for Samui

Samui supports image conversion from a TIFF file format. All TIFF images were transformed into the Cloud-optimized GeoTIFF format in Python 3.10 using GDAL v3.6.0 and rasterio v1.3.4. JPEG compression was performed using gdal_translate at quality 90. An example script to generate a COG from an example TIFF file is available at https://github.com/chaichontat/samui/blob/f4c886afe92400a651cbc59619b6060975b264c7/scripts/process_image.py. Overlays and feature data can be converted from a pandas DataFrame format using the provided Python API.

### Visium Spatial Gene Expression Data

Full-resolution images, filtered spaceranger output, and annotations were obtained from [26] (https://github.com/LieberInstitute/HumanPilot). The script to transform this data is available at https://github.com/chaichontat/samui/blob/f4c886afe92400a651cbc59619b6060975b264c7/scripts/process_humanpilot.py. Briefly, filtered outputs from 10x Genomics spaceranger v1.3.1 were opened using anndata v0.8 (https://anndata.readthedocs.io/en/latest) and log-transformed into a chunked CSV format. Images were Gaussian filtered with a radius of 4 to reduce artifacts, and processed into the Samui format as described above.

### MERFISH Data

Cell by gene data matrices were obtained from the Vizgen data release program [27, 28]. Images from the first *z*-stack were used as the background. The overlay is the position of each cell and features are the counts of MERFISH spots in each cell. Images and overlays were aligned using the provided affine transformation matrix. The script to transform Vizgen-formatted data is available at https://github.com/chaichontat/samui/blob/f4c886afe92400a651cbc59619b6060975b264c7/scripts/process_merfish.py.

### Visium Spatial Proteogenomics (Visium-SPG) Data

Full-resolution IF images, filtered spaceranger output, and annotations were obtained from [23] (https://github.com/LieberInstitute/spatialDLPFC). Primary antibodies were for four brain cell-type marker proteins, NeuN for neurons, TMEM119 for microglia, GFAP for astrocytes, and OLIG2 for oligodendrocytes.

## Abbreviations

AWS: amazon web services
COG: cloud-optimized GeoTIFF
CSV: comma-separated value
GIS: geographic Information systems
GUI: graphical user interface
HDF5: hierarchical data format version 5
SRT: spatially-resolved transcriptomics
TIFF: tag image file format

## Declarations

### Ethics approval and consent to participate

Not applicable.

### Consent for publication

Not applicable.

### Availability of data and materials

Samui is available at https://samuibrowser.com (v1.0.0-next.7). The 10x Genomics Visium data was obtained from [23, 26]. The source MERFISH data is available from the Vizgen Data Release Program [27, 28]. The code to reproduce the analyses and figures is available at https://github.com/chaichontat/samui/tree/f4c886afe92400a651cbc59619b6060975b264c7 and the software for the browser is available from https://github.com/chaichontat/samui.

### Author Contributions

- Conceptualization: CS, SCH
- Data Curation: CS
- Formal Analysis: CS, SCH
- Funding acquisition: KM, KRM, SCH, LCT
- Investigation: CS, AN, NJE, KM, KRM, SCH
- Project Administration: KM, KRM, SCH
- Software: CS, AN
- Supervision: KM, KRM, SCH
- Writing – original draft: CS, SCH
- Writing – review & editing: CS, AN, NJE, KM, KRM, SCH

## Acknowledgements

We would like to thank Sang Ho Kwon, Madhavi Tippani and Stephanie Cerceo Page for generation and processing of spatial transcriptomics images and data that were utilized to demonstrate the utility of Samui as well as for helpful comments, feedback and suggestions on Samui functionality.

## Bibliography

1. Marx V. Method of the Year: spatially resolved transcriptomics. Nat Methods. 2021;18:9–14.

2. Way GP, Spitzer H, Burnham P, Raj A, Theis F, Singh S, et al. Image-based profiling: a powerful and challenging new data type. Pac Symp Biocomput. 2022;27:407–11.

3. Schroeder AB, Dobson ETA, Rueden CT, Tomancak P, Jug F, Eliceiri KW. The ImageJ ecosystem: Open-source software for image visualization, processing, and analysis. Protein Sci. 2021;30:234–49.

4. Sofroniew N, Evans K, Nunez-Iglesias J, Solak AC, Lambert T, Kevinyamauchi, et al. napari/napari: napari 0.2.6. Zenodo. 2019.

5. Levet F, Carpenter AE, Eliceiri KW, Kreshuk A, Bankhead P, Haase R. Developing open-source software for bioimage analysis: opportunities and challenges. F1000Res. 2021;10:302.

6. GitHub - google/neuroglancer: WebGL-based viewer for volumetric data. https://github.com/google/neuroglancer. Accessed 17 Aug 2022.

7. Chang W, Cheng J, Allaire JJ, Sievert C, Schloerke B, Xie Y, et al. shiny: Web Application Framework for R. Computer software. NA; 2022. https://shiny.rstudio.com/. Accessed 20 Dec 2022.

8. OpenLayers - Welcome. https://openlayers.org/. Accessed 21 Dec 2022.

9. Cloud Optimized GeoTIFF. https://www.cogeo.org/. Accessed 21 Dec 2022.

10. Ståhl PL, Salmén F, Vickovic S, Lundmark A, Navarro JF, Magnusson J, et al. Visualization and analysis of gene expression in tissue sections by spatial transcriptomics. Science. 2016;353:78–82.

11. Spatial Gene Expression - 10x Genomics. https://www.10xgenomics.com/products/spatial-gene-expression. Accessed 21 Dec 2022.

12. Rodriques SG, Stickels RR, Goeva A, Martin CA, Murray E, Vanderburg CR, et al. Slide-seq: A scalable technology for measuring genome-wide expression at high spatial resolution. Science. 2019;363:1463–7.

13. Stickels RR, Murray E, Kumar P, Li J, Marshall JL, Di Bella DJ, et al. Highly sensitive spatial transcriptomics at near-cellular resolution with Slide-seqV2. Nat Biotechnol. 2021;39:313–9.

14. Eng C-HL, Lawson M, Zhu Q, Dries R, Koulena N, Takei Y, et al. Transcriptome-scale super-resolved imaging in tissues by RNA seqFISH. Nature. 2019;568:235–9.

15. Xia C, Fan J, Emanuel G, Hao J, Zhuang X. Spatial transcriptome profiling by MERFISH reveals subcellular RNA compartmentalization and cell cycle-dependent gene expression. Proc Natl Acad Sci USA. 2019;116:19490–9.

16. The HDF5^®^ Library & File Format - The HDF Group. http://www.hdfgroup.org/HDF5. Accessed 21 Dec 2022.

17. Zarr — zarr 2.13.3 documentation. https://zarr.readthedocs.io/en/stable/. Accessed 21 Dec 2022.

18. HTTP | MDN. https://developer.mozilla.org/en-US/docs/Web/HTTP. Accessed 21 Dec 2022.

19. OGC GeoTIFF Standard | OGC. https://www.ogc.org/standards/geotiff. Accessed 21 Dec 2022.

20. Nusrat S, Harbig T, Gehlenborg N. Tasks, techniques, and tools for genomic data visualization. Comput Graph Forum. 2019;38:781–805.

21. Leaflet - a JavaScript library for interactive maps. https://leafletjs.com/. Accessed 21 Dec 2022.

22. Diblen F, Hendriks L, Stienen B, Caron S, Bakhshi R, Attema J. Interactive Web-Based Visualization of Multidimensional Physical and Astronomical Data. Front Big Data. 2021;4:626998.

23. Huuki-Myers LA, Spangler A, Eagles NJ, Montgomery KD, Kwon SH, Guo B, et al. Integrated single cell and unsupervised spatial transcriptomic analysis defines molecular anatomy of the human dorsolateral prefrontal cortex. BioRxiv. 2023.

24. Dries R, Zhu Q, Dong R, Eng C-HL, Li H, Liu K, et al. Giotto: a toolbox for integrative analysis and visualization of spatial expression data. Genome Biol. 2021;22:78.

25. Pardo B, Spangler A, Weber LM, Page SC, Hicks SC, Jaffe AE, et al. spatialLIBD: an R/Bioconductor package to visualize spatially-resolved transcriptomics data. BMC Genomics. 2022;23:434.

26. Maynard KR, Collado-Torres L, Weber LM, Uytingco C, Barry BK, Williams SR, et al. Transcriptome-scale spatial gene expression in the human dorsolateral prefrontal cortex. Nat Neurosci. 2021;24:425–36.

27. Data Release Program | Vizgen. https://vizgen.com/data-release-program/. Accessed 21 Dec 2022.

28. Vizgen MERFISH Mouse Receptor Map. https://info.vizgen.com/mouse-brain-data. Accessed 10 Nov 2022.

